# Control of a Bi-Stable Genetic System via Parallelized Reinforcement Learning

**DOI:** 10.1101/2025.09.14.676149

**Authors:** Robin Henry, Jean-Baptiste Lugagne

**Affiliations:** Department of Engineering Science, University of Oxford, Oxford, United Kingdom

## Abstract

Achieving real-time control of genetic systems is critical for improving the reliability, efficiency, and reproducibility of biological research and engineering. Yet the intrinsic stochasticity of these systems makes this goal difficult. Prior efforts have faced three recurring challenges: (a) predictive models of gene expression dynamics are often inaccurate or unavailable, (b) nonlinear dynamics and feedback in genetic circuits frequently lead to multi-stability, limiting the effectiveness of deterministic control strategies, and (c) slow biological response times make data collection for learning-based methods prohibitively time-consuming. Recent experimental advances now allow the parallel observation and manipulation of over a million individual cells, opening the door to model-free, data-driven control strategies. Here we investigate the use of Parallelized Q-Networks (PQN), a recently-developed reinforcement learning algorithm, to learn control policies for a simulated bi-stable gene regulatory network. We show that PQN can not only control this self-activating system more accurately than other model-free and model-based control methods previously used in the field, but also converges efficiently enough to be practical for experimental application. Our results suggest that parallelized experiments coupled with advances in reinforcement learning provide a viable path for real-time, model-free control of complex, multi-stable biological systems.

## I. Introduction

The ability to precisely manipulate biological systems could transform biomanufacturing, healthcare, and agriculture. Effective gene expression control is central to this effort, yet remains difficult due to the stochastic, nonlinear, and often poorly characterized dynamics of gene regulation networks [1], [2].

The emerging field of cybergenetics has made significant progress toward addressing these challenges through realtime feedback control of gene expression at the single-cell level [3]. Such approaches typically fall into two categories: model-free controllers like bang-bang or proportional integral (PI) [4], [5], which require minimal prior system knowledge but often lack accuracy, and model-based methods like model predictive control (MPC) [6], [7], which rely heavily on expert-derived nonlinear ordinary differential equation (ODE) models. While model-based approaches often perform better, their dependence on accurate models limits applicability in uncertain systems such as gene regulation networks.

Recent advances in microscopy automation, image analysis, and microfluidics now allow parallel observation and manipulation of up to one million cells [8]. Leveraging this, we previously introduced deep model predictive control (deep MPC), which uses a machine learning model within an MPC framework, to control gene expression across thousands of bacterial cells. This enabled unprecedented accuracy and throughput in real-time control and helped uncover dynamics of antibiotic resistance in bacteria [9], [10].

Here, we explore whether recent advances in deep reinforcement learning (RL), particularly those designed for massive parallelization, can further improve control of complex biological systems without prior knowledge of their dynamics. We focus on multi-stable genetic circuits — common in processes like bacterial decision-making and differentiation [11] — and evaluate control strategies on a self-activating gene circuit exhibiting inherent bi-stability driven by a positive feedback loop. This simple circuit is coupled to a light-based, or optogenetic, activation system that we used and studied previously [12], [9], [10]. We hypothesized that deep MPC, which predicts a single likely trajectory, would be suboptimal for such systems, while RL’s capacity for long-term decision-making might help avoid suboptimal attractor states.

RL algorithms, such as Proximal Policy Optimization (PPO) [13], are often considered sample inefficient, and various strategies have been proposed to mitigate the impact of the slow timescale of biological systems like sim-to-real approaches [14]. But biology offers a unique advantage: the ability to massively parallelize experiments [8]. This could overcome traditional efficiency barriers by enabling thousands of simultaneous learning interactions. Motivated by recent introduction of Parallelized Q-Networks (PQN) [15], a Deep Q-Network (DQN) variant that trains across a large number of environments in parallel, we explore whether PQN can be paired with high-throughput experimental platforms to rapidly learn effective control policies for systems with unknown dynamics.

In this study, we systematically compare bang-bang, PI, and deep MPC against RL-based PPO and PQN approaches on our simulated bi-stable circuit. Our results show that PQN outperforms all other methods in control accuracy while converging fast enough for real-time experimental application. These findings suggest that combining deep RL with parallelized experiments offers a powerful new approach for controlling complex biological systems without a priori models.

## II. Preliminaries

This section introduces key concepts, notations, and terminologies used throughout the rest of this paper.

### A. Reinforcement Learning

In the RL literature, control tasks are modeled as Markov Decision Processes (MDP), where an agent interacts with an environment over an infinite sequence of discrete time steps {0, 1, … } [16]. At each time step *t* ∈ 𝒯, the agent selects an action *a*_*t*_ ∈ 𝒜 based on a state *s*_*t*_ ∈ 𝒮 according to a stochastic policy *π* : *S* × *A* → [0, 1], such that *a*_*t*_ ∼ *π*(· | *s*_*t*_). After the action is applied, the agent transitions to a new state *s*_*t*+1_ ∼ *p*(· | *s*_*t*_, *a*_*t*_) ∈ 𝒮 and receives a reward *r*_*t*_ := *r*(*s*_*t*_, *a*_*t*_, *s*_*t*+1_) ∈ ℝ. As the agent interacts with the environment, it collects a trajectory *τ*_*t*_ := (*s*_0_, *a*_0_, *r*_0_, *s*_1_, …, *s*_*t*−1_, *a*_*t*−1_, *r*_*t*−1_, *s*_*t*_) with distribution 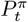. The expected return *J*^*π*^ associated with policy *π* is defined as:

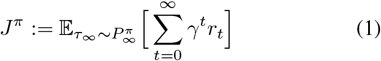

where *γ* ∈ [0, 1) is the discount factor enforcing convergence. The agent’s goal is to learn an optimal policy *π*^*^ that maximizes its expected discounted return 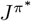.

In partially-observable environments, such as the one considered in this paper, the agent only has access to observations *o*_*t*_ ∈ 𝒪 and not to the full state *s*_*t*_. In such cases, the agent must infer, directly or indirectly, a representation of the state from the observable trajectory (*o*_0_, *a*_0_, *r*_0_, …, *o*_*t*−1_, *a*_*t*−1_, *r*_*t*−1_, *o*_*t*_) [17].

### B. Gillespie Algorithm

Efficient computational simulations of a set of biochemical reactions within cells is often done via the Gillespie algorithm [18], a Monte Carlo method also known as the Stochastic Simulation Algorithm [19].

Consider a set of *m* chemical species (e.g., proteins) *x*_*i*_, ∀*i* ∈ {1, …, *m*}, denoted by the vector x ∈ ℤ^*m*^, and a set of *n* possible reactions acting on those species, each described by a propensity function *p*_*i*_ : ℤ^*m*^ → ℝ^+^, ∀*i* ∈ {1, …, *n* }, such that *p*_*i*_(x) denotes the propensity of reaction *i* given the current species population x. Let v_*i*_ ∈ ℤ^*m*^ be the stoichiometry vector describing the change in species counts that occurs when reaction *i* fires. The Gillespie algorithm simulates trajectories of this system by iterating over the following steps until some termination criterion is reached.

*1) Time step sampling:* Sample the time lapse until the next reaction starts, *δt*, from an exponential distribution with mean 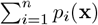:

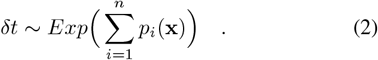

*2) Reaction selection:* Choose the reaction *j* to simulate by sampling *j* ∼ *P* (· | x), where the probability of reaction *i* happening next is:

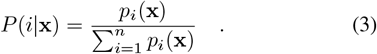

*3) State update:* Update the population x according to the stoichiometry of reaction *j*, and increment time by *δt*:

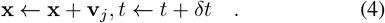

In our case, the simulation is terminated once time *t* reaches a predetermined sampling time used by our control agent.

## III. Control Task

This section introduces the optogenetic system used throughout this paper and formulates the control task we aim to solve as an MDP.

### A. Biological System

We focused our efforts on a simulated optogenetic circuit with a self-activating loop that yields bistable dynamics. Through a cascade of biochemical reactions, the production of a green fluorescent protein (GFP, our measured output) is activated under green light; red light deactivates GFP expression, after which cell fluorescence decays to zero due to dilution and cell division (Fig. 1A). Experimentally, singlecell light stimulation and fluorescence readout are made possible by specialized microscopy [9].

**Fig. 1.**
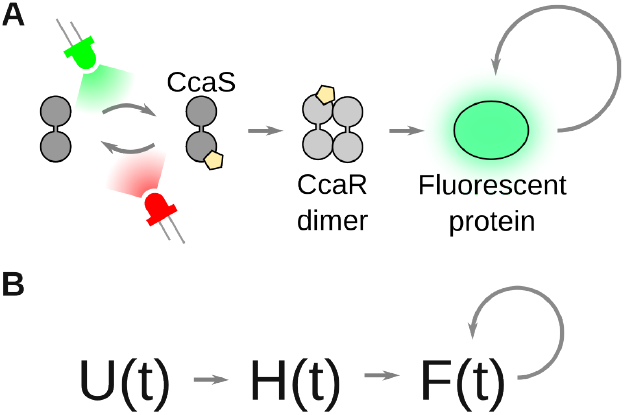
The self-activating optogenetic system we aim to control. A) Green light activates CcaS, which catalyzes the formation of CcaR dimers that drive expression of a GFP reporter protein, which further activates itself. Red light deactivates CcaS, leading to GFP proteins diluting due to growth and division. B) The simplified circuit we simulated: a light input signal *U* (*t*) modulates the formation of *H*(*t*) dimers, which activate the expression of *F* (*t*) proteins that also self-activate.

More specifically, we modeled the CcaS-CcaR system [12]: green light activates the light-sensitive CcaS protein, which in turn activates the CcaR protein, leading to the formation of CcaR dimers. These dimers then activate the transcription of a reporter, such as GFP. Conversely, red light deactivates CcaS and the system returns to a low-GFP state. We further added positive feedback by letting GFP activate its own expression while noting that, in practice, this would require a co-expressed transcription factor or indirect means like growth inhibition, which we chose to omit for simplicity.

We modeled a simplified version of this system, as shown in Fig. 1B. *U* (*t*) ∈ {0, 1} denotes the time-varying light input at time step *t* (0 = red, 1 = green), *H* ∈ (*t*) ℤ encapsulates the dynamics of the optogenetic cascade and represents activated CcaR proteins, and *F* (*t*) ∈ ℤ is the number of fluorescent reporter proteins. The reaction stoichiometries and corresponding propensity functions for the CcaSR self-activating system are summarized in Table I, where:

- *ν* = 0.01 is the dilution rate of proteins,
- *K*_*H*_ = 90 and *n*_*H*_ = 3.6 are the Hill function parameters for the activation of F proteins from H, and
- *K*_*F*_ = 30 and *n*_*F*_ = 3.6 are the Hill function parameters for the self-activation of F proteins.

**TABLE I.** Gillespie reactions and propensities for the CcaSR self-activating system.

| Reaction $i$ | Propensity $p_i(\mathbf{x})$ |
| --- | --- |
| $\emptyset \rightarrow H$ | $U$ |
| $H \rightarrow \emptyset$ | $\nu H$ |
| $\emptyset \rightarrow F$ | $\frac{1}{2} \frac{H^{n_H}}{K_H^{n_H} + H^{n_H}}$ |
| $\emptyset \rightarrow F$ | $\frac{1}{2} \frac{F^{n_F}}{K_F^{n_F} + F^{n_F}}$ |
| $F \rightarrow \emptyset$ | $\nu F$ |

Parameters are normalized from experimentally-derived values, taken from [20].

### B. System Analysis

To perform a bifurcation analysis of the self-activating gene regulatory system, we first established a deterministic, continuous-time ODE model. The ODEs were derived from the reaction propensities detailed in Table I and are described by the following equations:

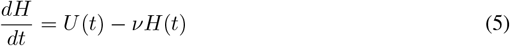

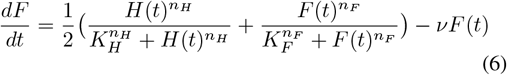

We identified the equilibrium points of the system for constant input values *U* (*t*) = *u* ∈ [0, 1]. These constant inputs can be viewed as averaged approximations of rapidly alternating input signals between red and green illumination. The resulting bifurcation diagram in Fig. 2A demonstrates a clear hysteresis cycle in the steady-state values of *F* (*t*). Specifically, the system exhibits two stable steady states and an intermediate unstable equilibrium for 0.42 *< u <* 0.64. Therefore, dynamic control is necessary to stabilize fluorescence output in the intermediate unstable region 16.2 *< F* (*t*) *<* 38.1.

**Fig. 2.**
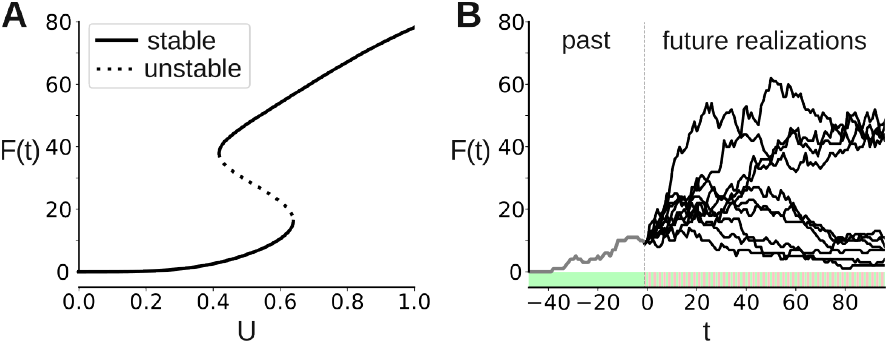
The self-activating optogenetic system is bi-stable. A) Bifurcation diagram of the deterministic model. B) Diverging realizations of stochastic fluorescence responses to the same sequence of alternating red and green light inputs.

To further illustrate the implications of bi-stability, we ran a stochastic Gillespie simulation of our system. All molecular species were initialized to zero, and the simulated cell was subjected initially to continuous green illumination (*U* (*t*) = 1) for 48 time points (4 in-simulation hours) to activate gene expression to an intermediate value. Then, the cell was exposed to an alternating sequence of red and green light (*U* (*t*) = {0, 1, 0, 1, … }). We obtained 10 independent stochastic realizations of this simulation, and show the resulting fluorescence trajectories over time in Fig. 2B. Four out of ten simulated trajectories converged to the higher basin of attraction, while the others remained at lower levels of *F* (*t*).

These results highlight the inherent difficulty of controlling stochastic multi-stable genetic systems. Identical control strategies can lead to divergent outcomes, and early decisions can have long-term consequences that create persistent error.

### C. MDP Task

In the most general sense, we were interested in studying to what extent various algorithms are able to control the number of F proteins over time without expert knowledge of the system. In other words, whether they can follow some objective function 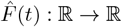.

At time *t*, our system is fully characterized by the number of molecules of each species. Hence, 𝒮 := ℤ^2^, with *s*_*t*_ := [*H*(*t*), *F* (*t*)].

However, during practical experiments, the system is partially observable: we only measure fluorescence intensity, a proxy for *F* (*t*). For simplicity, we make the assumption here that a perfect mapping from fluorescence intensity to *F* (*t*) is available, i.e. that the agent observes the true *F* (*t*). Rather than providing perfect knowledge of 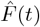 to the controller, we also restrict its access to the sequence of future 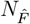 objective values. The agent thus receives observation vectors of the form:

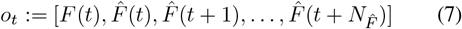

with 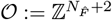.

Actions are boolean values corresponding to red (0) or green (1) light, so 𝒜 := *{*0, 1*}*.

We emulated state transitions by running stochastic simulations of our genetic circuit. Given a starting state *s*_*t*_ and an action *a*_*t*_, the next state *s*_*t*+1_ is obtained by generating a trajectory of the cell species via the Gillespie algorithm (see Section II-B) over the continuous interval [*t, t* + 1). During these Gillespie simulations, the light input value *U* used in propensity equation *p*_1_ (see Table I) is kept constant at the action value *a*_*t*_.

Transition rewards are calculated as the negative absolute distance between the current number of reporter proteins *F* (*t*) and the target 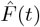:

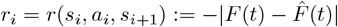

It is worth noting that this formulation is identical to RMSE with one-element vectors.

## IV. Control Algorithms

Since we were interested in controlling biological systems with unknown dynamics, we tested a series of control algorithms that do not require a priori expert knowledge of the system. Namely, we tested random, bang-bang, and PI controllers, along with deep MPC and RL-based PPO and PQN algorithms.

### A. Baselines

We ran all our experiments with three baseline algorithms of increasing complexity: a random policy, a bang-bang controller, and a PI controller.

At time *t*, the random policy chooses a random action 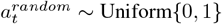.

The bang-bang controller follows the policy:

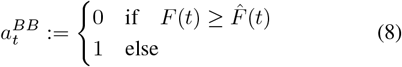

in an attempt to move *F* (*t*) closer to the next 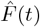.

For the PI controller, let *u*(*t*) be the continuous PI control signal computed at time *t*:

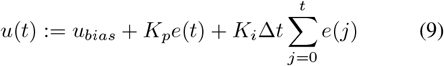

where 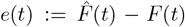 is the current error term, Δ*t* is the sampling time, *K*_*p*_ and *K*_*i*_ are the proportional and integral gains, respectively, and *u*_*bias*_ is a constant bias. The final binary action is given by the unit step function:

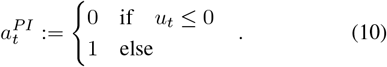

### B. Deep Model Predictive Control

We hypothesized that forward-looking methods would perform better than the baselines. For this reason, and because it performed well in our previous study [9], we next tested a deep MPC algorithm.

Deep MPC is an MPC variant that uses a deep neural network for the prediction step. MPC involves, at each time step, predicting the future evolution of a system’s state using a mathematical model and then optimizing a sequence of control actions over a finite prediction horizon [21]. The neural network we used for predictions featured a long-short term memory (LSTM)-based encoder followed by a fully connected neural network (MLP) decoder. We trained it to predict future fluorescence responses given (a) past optogenetic inputs, (b) the fluorescence history, and (c) the future optogenetic input sequence. This was done by randomly splitting simulated trajectories into past and future windows and minimizing the mean squared error loss. Our training dataset consisted of 10,000 input-output pairs of 432 time points (equivalent to 36 hours) where random input sequences were generated by thresholding a bounded one-dimensional random walk as described in [4].

To control gene expression, we then integrated the trained neural network model into an MPC framework, using a binary particle swarm optimizer [22] to find near-optimal control strategies. The objective of the optimizer was to minimize the root mean squared error between the predicted cell responses and a specified control objective trajectory over the prediction horizon. The MPC control policy can be expressed as:

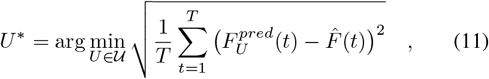

where *U* ^*^ is the optimal control input sequence, *T* is the prediction horizon, 𝒰:= {0, 1}^*T*^ is the set of candidate binary sequences of length *T*, and 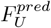 represents the fluorescence response predicted by the neural network under optogenetic sequence *U* .

The only difference in implementation between this study and our previous work [9] is an extension of the prediction horizon from *T* = 24 time points (equivalent to 2 hours with Δ*t* = 5 minutes) to *T* = 96 time points (8 hours) to prevent issues where short-horizon planning resulted in simulated cells getting trapped in stable states.

### C. Proximal Policy Optimization

Early on, we hypothesized that deep MPC might perform poorly when applied to the bi-stable genetic system studied here due to the neural network predicting only a single, “most likely” trajectory. Approaches relying on such deterministic predictions are inherently limited for systems with multimodal responses, as it does not adequately capture the probabilistic dynamics of our bi-stable self-activating genetic loop. In contrast, RL-based methods are free from this limitation.

As a stable and effective on-policy policy gradient RL algorithm, PPO [13] is often the go-to option for RL tasks. It alternates between collecting experience, in the form of finite-length trajectories following the current policy, and performing several epochs of optimization on the collected data to update the current policy, after which the collected experience is discarded. During each policy update step, the policy parameters are updated by maximizing a clipped objective function characterized by a hyperparameter that dictates how far away the new policy is allowed to diverge from the old one. The objective also requires the use of an advantage-function estimator, which is achieved using a learned-state value function.

Although PPO has been successfully deployed on a wide range of RL tasks due to its stability w.r.t. hyperparameter choice, it remains sample inefficient, as illustrated by our results in Section V-C.

### D. Parallelized Q-Networks

The hypothesis at the origin of this study was that PQN [15] would perform at least as well as the other algorithms while also drastically reducing the laboratory experimental time needed.

Q-learning is a model-free, off-policy algorithm. It aims to learn an action-value function *Q* : 𝒮 *×* 𝒜 → ℝ such that *Q*(*s, a*) estimates the total future rewards from state *s* if action *a* is taken. Enacting the policy learned is then as simple as always taking the action with the largest Q-value, in a greedy manner:

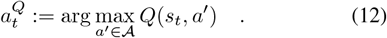

PQN is a simplified version of Deep Q-learning (DQN), which is itself a variant of Q-learning where the Q-function is approximated by a neural network, often referred to as the Q-network. Notably, PQN removes the need of large replay buffers during training by taking advantage of many parallel environments. We hypothesized that experimental capabilities in growing and observing large numbers of cells (i.e., environments) in parallel would enable it to learn more quickly than other RL algorithms such as PPO.

We experimented with two different architectures for the Q-network of PQN agents, inspired by [15]. The first was a simple MLP network, where we stacked and flattened the history of the last *N*_*obs*_ observation vectors as input to the network, as is common when training agents with memory-less networks in partially observable environments. The second architecture had an additional recurrent neural network (RNN) component, to encode a memory of previous observations. Because we consistently reached better performance (although by thin margins) and faster convergence speed with the MLP variant, we chose to omit results for the RNN version, for brevity. However, biological systems with longer-term dependencies might benefit from the use of an RNN encoder.

## V. Results

### A. Experimental Setup

The results presented here focus on a particular instance of the control task introduced in Section III, in which the goal is to keep the number of GFP proteins *F* (*t*) within the unstable region. More specifically, we set the objective function to a constant 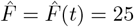, ∀*t*, and only provided 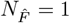 future objective values to optimize memory usage, since a larger value would simply add duplicate values to the observation vector *o*_*t*_ without providing the agent with more information. We used an in-simulation sampling time Δ*t* of 5 minutes, as is common in experimental setups, and limited episode lengths to 24 hours, making each trajectory 288 discrete time steps long. To remain consistent with laboratory conditions where cells are exposed to constant red light before activating control, all episodes started from a fixed initial state *s*_0_ = [0, 0].

Inspired by previous works [23], our experimental framework was implemented primarily in JAX [24], taking advantage of just-in-time compilation and highly parallelized end-to-end training and evaluation of control and RL algorithms. We implemented the control task using the gymnax framework [25] and our PQN implementations were highly inspired by the purejaxrl repository released with [15]. We ran Bayesian hyperparameter optimization sweeps using the wandb framework [26] for the PI controller and all RL models on single-thread, single-GPU nodes of the University of Oxford HPC cluster. Our code is available at https://gitlab.com/lugagnelab/pqn-control-cdc2025.

The final performance of each algorithm was evaluated by reporting the mean and standard deviation of the total sum of rewards (non-discounted return) over 1,000 288-step-long episodes and 3 random seeds. For RL algorithms, we also evaluated performance throughout training in order to assess policy convergence speed and stability. Every *N*_*updates*_*/*100 policy update steps, where *N*_*updates*_ refers to the total number of update steps, the agents were evaluated on 500 episodes.

### B. Control Accuracy

The first question we wanted to answer was whether the RL-based approaches would perform at least as well as the baselines and deep MPC methods. The answer is yes, as illustrated in Fig. 3 and summarized in Table II.

**TABLE II.** Mean and standard deviation of episode returns for each algorithm, evaluated over 1,000 episodes and 3 random seeds. The middle column contains statistics for the whole 24-hour period, whereas the right column only considers hours 15 to 24, past the initial transient period.

| Algorithm | Return (hours 0-24) | Return (hours 15-24) |
| --- | --- | --- |
| Random | -5,905 ± 496 | -780 ± 150 |
| Bang-Bang | -3,028 ± 770 | -365 ± 219 |
| PI | -2,914 ± 682 | -365 ± 198 |
| Deep MPC | -2,821 ± 732 | -334 ± 227 |
| PPO | -2,605 ± 750 | -311 ± 224 |
| **PQN** | **-2,455 ± 633** | **-281 ± 183** |

**Fig. 3.**
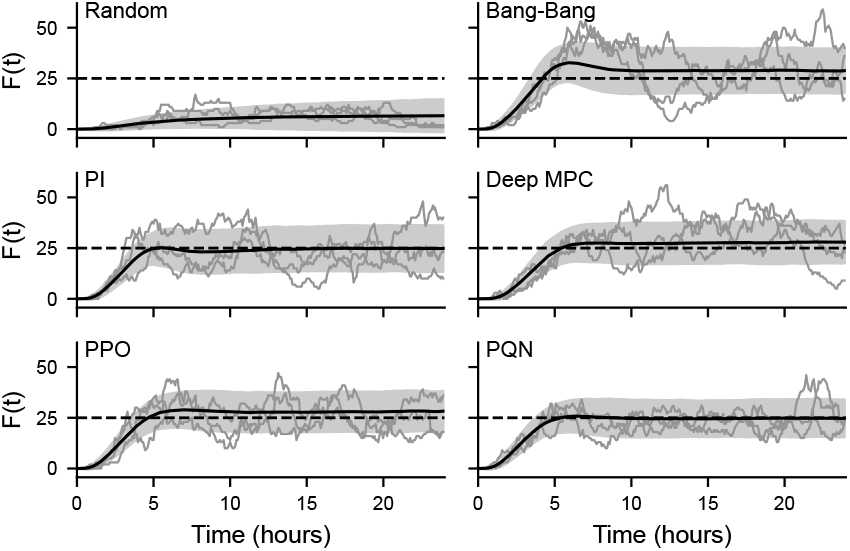
Trajectories of the number of GFP proteins, *F* (*t*), over time, for different control algorithms. Each plot contains 3 randomly selected individual trajectories (gray lines), the average trajectory (solid black line) with its standard deviation (shaded area), and the target 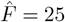 (dashed line). The mean and standard deviations were obtained from 1,000 24-hour episodes and 3 random seeds.

As expected, the random policy is trapped in the lower basin of attraction, leading to large negative returns (Table II). The bang-bang approach performs better but, on average, largely overshoots early on and fails to recover, settling around *F* (*t*) = 28 (Fig. 3). Thanks to its integral component, the PI controller manages to track 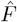 more accurately on average, but tends to induce a longer settling time. In addition, trajectories from both the bang-bang and PI controllers suffer from large standard deviations, as illustrated by the trajectory examples of Fig. 3. On average, the deep MPC model outperforms all three baseline algorithms. Both RL algorithms reach better performance however, with PQN yielding the best results, earning the highest average return and lowest standard deviation of all algorithms tested (Table II).

A large proportion of the total return of trajectories can be attributed to the first 5 hours, during which all algorithms need time to get out of the lower initial state and closer to 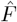 (see Fig. 3). In addition, the bang-bang and PI controllers produce long transient responses, stabilizing around hours 10 and 15, respectively. To assess the steady-state performance of the different methods, Table II also summarizes return scores over hours 15 to 24 (right column). Zooming in on those hours increases the gap between the baseline algorithms and the deep MPC model: from 3.2% to 8.5% between the PI controller and deep MPC. However, the RL approaches remain the best performing algorithm, scoring 6.9% (PPO) and 15.9% (PQN) increases in average return over deep MPC.

### C. Training Convergence

The second question we were interested in was whether the RL policies would converge within a timeframe reasonable for practical experiments. Results are shown in Fig. 4 and Table III. PPO performance improves relatively quickly in the first 20 hours before progressing incrementally, while PQN converges significantly faster overall. This validates our early hypothesis that the parallelization capabilities of PQN would make it a more viable approach.

**Fig. 4.**
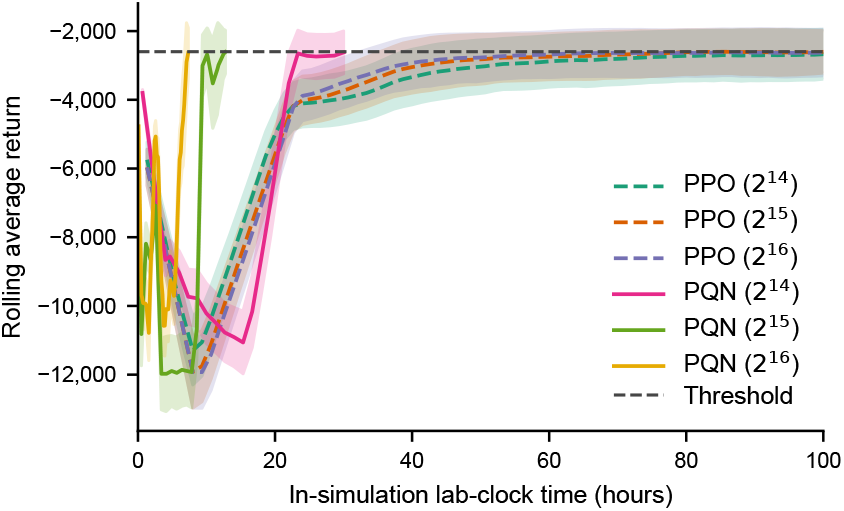
Rolling performance (mean episodic return) of PPO and PQN agents during training, averaged over 500 episodes and 3 random seeds.

Specifically, we evaluated the effect of increasing the number of simulated cells on which the agent is trained in parallel from 2^10^ = 1, 024 cells to 2^16^ = 65, 536 cells. We selected an average return of -2,600 as the threshold to reach for an agent to be considered fully trained, shown as a horizontal line in Fig. 4. We chose this value as a middle ground between the -2,821 and -2,455 average return reached by deep MPC and our best PQN agents, respectively (see Table II).

Both algorithms exhibit an inverse relationship between the number of parallel environments used for training and the convergence speed (in terms of simulated laboratory-clock time), as seen in Table III. The only notable exception being from 2^15^ to 2^16^ cells for PPO, where the runtime starts to slightly increase again.

**TABLE III.** The equivalent laboratory experimental time needed for PPO and PQN agents to reach an average return of -2,600, for different number of parallel environments.

| # Environments | PPO (hours) | PQN (hours) |
| --- | --- | --- |
| 1,024 ($2^{10}$) | 2,158.7 | 317.3 |
| 8,192 ($2^{13}$) | 169.3 | 34.7 |
| 16,384 ($2^{14}$) | 128.0 | 30.7 |
| 32,768 ($2^{15}$) | 102.7 | 12.8 |
| 65,536 ($2^{16}$) | 110.7 | **7.4** |

PQN clearly outperforms PPO in terms of convergence speed, reaching as low as 7.4 hours with 65k (2^16^) environments against *>*100 hours for PPO. This is particularly impressive considering the 4 to 5 hours required for the genetic system to first reach *F* (*t*) = 25. While, due to GPU memory limitations, we were not able to investigate larger numbers of parallel environments, these results suggest that convergence time could be reduced even further in practice since the latest experimental platforms can operate on up to one million cells in parallel [8].

## VI. Conclusion

Overall, our results suggest that interfacing RL agents with living cells is feasible and promising for controlling biological systems. By combining Parallelized Q-Networks with emerging high-throughput experimental platforms for singlecell control, we showed that RL can outperform state-ofthe-art methods in real-time gene expression control. Critically, our results show that the training time is practically feasible: PQN agents outperform deep MPC under 8 hours of simulated experimental time with 65,536 parallel cells, a duration that is likely to decrease further with advances in experimental throughput [27], [8].

Our current simulations do not account for factors like cell-to-cell variability, long-term dynamics such as cell aging, and measurement noise, all of which complicate gene expression control. Future research will refine our simulation environment to better reflect experimental realities. Additionally, we plan to explore other gene networks and RL algorithms to assess the approach’s generalizability and hyperparameter robustness.

Still, based on our prior work with similar models, this study offers valuable insights into the potential of RL to manipulate complex biological systems, and into the data requirements for practical laboratory implementation.

## Acknowledgment

The authors thank Matteo Gallici and Mattie Fellows for insightful discussions regarding PQN implementation.

The authors would also like to acknowledge the use of the University of Oxford Advanced Research Computing (ARC) facility in carrying out this work. http://dx.doi.org/10.5281/zenodo.22558

RH was supported by the Engineering and Physical Sciences Research Council (EPSRC) UK, Centre for Doctoral Training in Engineering Biology (grant EP/Y034791/1) and by the EPSRC EEBio Programme (grant EP/Y014073/1).

